# Validation of selective agars for detection and quantification of *Escherichia coli* resistant to critically important antimicrobials

**DOI:** 10.1101/2021.06.18.449085

**Authors:** Zheng Z. Lee, Rebecca Abraham, Mark O’Dea, Ali Harb, Kelly Hunt, Terence Lee, Sam Abraham, David Jordan

**Affiliations:** Antimicrobial Resistance and Infectious Diseases Laboratory, Murdoch University, Murdoch, WA, Australia; New South Wales Department of Primary Industries, Wollongbar, NSW, Australia

## Abstract

Success in the global fight against antimicrobial resistance (AMR) is likely to improve if surveillance can be performed more rapidly, affordably and on a larger scale. An approach based on robotics and agars incorporated with antimicrobials has enormous potential to achieve this. However, there is a need to identify the combinations of selective agars and key antimicrobials yielding the most accurate counts of susceptible and resistant organisms. A series of designed experiments involving 1,202 plates identified the best candidate-combinations from six commercially available agars and five antimicrobials using 18 *Escherichia coli* strains as either pure cultures or inoculums within faeces. The effect of various design factors on colony counts were analysed in generalised linear models. Without antimicrobials, Brilliance^™^ *E. coli* (Brilliance) and CHROMagar^™^ ECC (CHROMagar) agars yielded 28.9% and 23.5% more colonies than MacConkey agar. The order of superiority of agars remained unchanged when faecal samples with and without spiking of resistant *E. coli* were inoculated onto agars with or without specific antimicrobials. When incorporating antimicrobials at varying concentrations, it was revealed that ampicillin, tetracycline and ciprofloxacin are suitable for incorporation into Brilliance and CHROMagar agars at all defined concentrations. Gentamicin was only suitable for incorporation at 8 and 16 μg/mL while ceftiofur was only suitable at 1 μg/mL. CHROMagar^™^ ESBL agar supported growth of a wider diversity of extended-spectrum cephalosporin-resistant *E. coli*. The findings demonstrate the potential for combining robotics with agars to deliver AMR surveillance on a vast scale with greater sensitivity of detection and strategic relevance.

**IMPORTANCE:** Established models of surveillance for AMR in livestock typically have a low sampling intensity which creates a tremendous barrier to understanding the variation of resistance amongst animal and food enterprises. However, developments in laboratory robotics now make it possible to rapidly and affordably process high volumes of samples. Combined with modern selective agars incorporating antimicrobials, this forms the basis of a novel surveillance process for identifying resistant bacteria by chromogenic reaction including accurately detecting and quantifying their presence even when present at low concentration. As *Escherichia coli* is a widely preferred indicator bacterium for AMR surveillance, this study identifies the optimal selective agar for quantifying resistant *E. coli* by assessing the growth performance on agars with antimicrobials. The findings are the first step towards exploiting laboratory robotics in an up-scaled approach to AMR surveillance in livestock with wider adaptations in food, clinical microbiology and public health.

## INTRODUCTION

Antimicrobial resistance (AMR) has been identified as one of the most serious threats to animal and human health in the current era (1). A key component for controlling AMR is the conduct of surveillance to inform on the prevalence and spread of resistant bacteria. The livestock sector has become a focus for surveillance because of the potential for AMR to transfer to humans along the food chain. Food products with a propensity to be contaminated with animal microflora such as ground meat are increasingly included in surveillance because of the risk of zoonotic pathogens undergoing selection for resistance in the animal gut or acquiring resistance via horizontal gene transfer (2-5). In both food and livestock, commensals such as *Escherichia coli* have been widely exploited for use in AMR surveillance since they readily develop resistance during *in-vivo* exposure to antimicrobials and are easily isolated as a ubiquitous component of the gut microflora (6). A barrier for improving surveillance in food and livestock is that the microbroth dilution technique for evaluating antimicrobial susceptibility of bacterial colonies, as recommended by international reference organisations, are expensive and labour intensive though the process has adapted well to a clinical context (7, 8). In national surveillance programs, sampling must typically be constrained due to the aforementioned drawbacks of the microbroth dilution technique. For example, fewer than 300 commensal *E. coli* isolates are obtained from the same number of herds or flocks of a given animal species in a year with food product surveys similarly affected (9). The inferences that can be drawn from surveillance results are thus often constrained in scope and frequently fail to support decision making at the coalface of animal and food production where changes to production management to control AMR arguably stands to have the greatest benefit. Therefore, an enhanced approach is needed that can affordably assess a substantially larger number of isolates and samples within an authoritative design to produce evidence on an epidemiological rather than clinical scale.

The problem of scale described above has the potential to be resolved using advancements in laboratory robotics to achieve a vast improvement in throughput of antimicrobial sensitivity assays. Robotics impart the capacity to rapidly handle samples, bacterial colonies and broth cultures in a coordinated way that minimises errors that arise from manual processes (10). This opens the door for efficient detection and quantification of AMR in commensal *E. coli* by estimating the number of colonies resistant in a given volume of faeces or food. These data offer a robust alternative for decision making on AMR control measures in food-producing enterprises. Additionally, the computerised reading, reporting and delivery of results means decisions can be made sooner. One way that robotics can be exploited is through large-scale enumeration of resistant *E. coli* from food or faecal samples using a process akin to agar dilution technique for antimicrobial susceptibility testing (AST). Here, automated plating of diluted samples onto agars incorporated with antimicrobials is the foundation. However, conventional solid agar used for this form of AST such as Mueller-Hinton agar (MHA) or traditional selective agar such as MacConkey (MAC) agar are unsuitable because they make it impossible to identify the target bacteria based solely on colony morphology, and especially in the case of MHA, the growth of non-target bacteria is not adequately suppressed.

Fortunately, modern selective agars are now commercially available for isolating *E. coli*. These agars suppress most non-target organisms and achieve accurate colony identification using a chromogenic reaction (11). One key issue in the use of these agars is whether or not the activity of antimicrobials that are incorporated is compromised by other agar components leading to inaccurate counts of resistant *E. coli*. A second key issue is whether or not the plating of diluted samples containing all of the microbial genera that are naturally occurring in the original samples (faeces or food) interferes with the AST of the target organism (in this case commensal *E. coli*). Previous studies have shown that MAC agar incorporated with ciprofloxacin is able to selectively isolate ciprofloxacin-resistant *E. coli* (12-14), although tetracycline cannot be used with MAC in this way due to interference in antimicrobial activity by divalent cations (calcium and magnesium salts are an integral component of MAC agar) (15-19). Similarly, there is a need to evaluate the suitability of commercial selective agars targeting extended-spectrum cephalosporin (ESC)-resistant *E. coli* such as Brilliance^™^ ESBL (Brilliance ESBL) and CHROMagar^™^ ESBL (CHROMagar ESBL) agars for detection and enumeration under the same conditions. This study aims to address these issues through three objectives. The first is to identify which selective agars (Brilliance^™^ *E. coli* [Brilliance] and CHROMagar^™^ ECC [CHROMagar] agars) have the best *E. coli* growth performance (highest colony counts per agar when plated with standardised inoculum) for accurate enumeration of *E. coli* colonies. Secondly, to identify which combination of specific antimicrobial (ampicillin, tetracycline, gentamicin, ciprofloxacin and ceftiofur) concentrations of antimicrobial and selective agars achieve the most accurate enumeration of resistant *E. coli* (this includes equivalent evaluation of commercial agars for isolation of ESC-resistant *E. coli*). Thirdly, to assess whether the ability to detect and quantify resistance is reduced when the target organisms are co-mingled with natural flora present in faecal samples. Together, the findings will serve to identify the optimal selective agar for achieving large-scale detection and quantification of resistant *E. coli* in samples from the food chain using laboratory robotics.

## RESULTS

### Experiment A: Comparison of *E. coli* growth on commercial *E. coli* selective agars

Three selective agars and one non-selective agar without incorporation of antimicrobials were compared for the ability to support growth of diverse *E. coli* strains (with and without resistance to various antimicrobials: Table S1). All *E. coli* strains grew on agar without antimicrobials. Agar had a highly significant effect (P<0.01) on colony growth with the order of superiority being Brilliance agar (mean of 78.9 colonies per plate), CHROMagar agar (mean of 74.7 colonies per plate), MHA (mean of 60 colonies per plate) and MAC agar (mean of 59 colonies per plate) (Fig. 1). However, though strain did have a significant effect on colony counts (P<0.001), it did not change the above order of superiority of agars for any strain (Fig. S1). In summary, *E. coli* counts on Brilliance, CHROMagar and MHA were on average 28.9%, 23.5% and 1.68% respectively higher than MAC agar (the worst performing).

**FIG 1.**
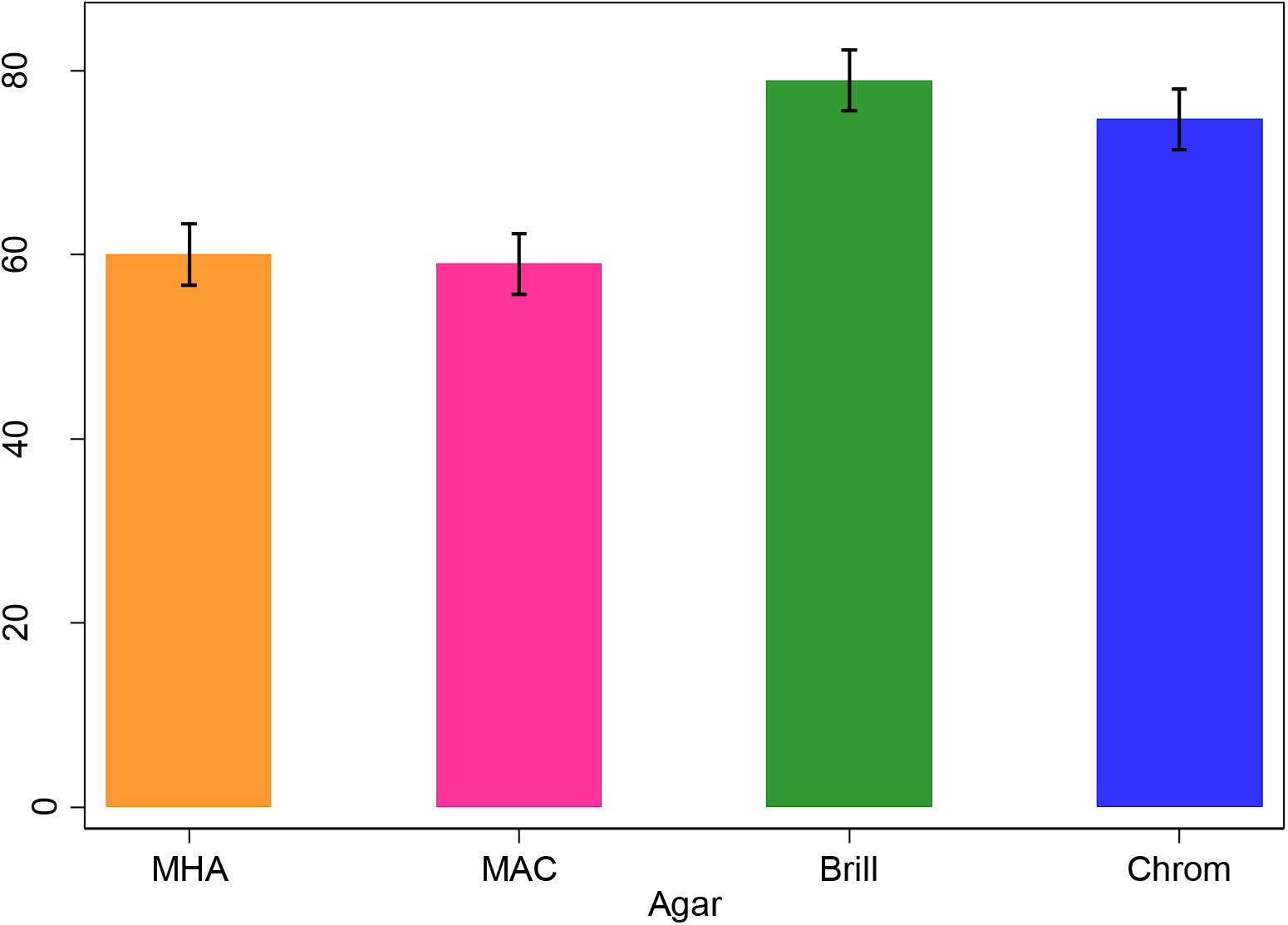
Comparisons of *E. coli* growth performance (mean colony counts per plate ± se) on three *E. coli* selective agar and Mueller-Hinton agar (all without antimicrobials) (total number of plates = 160). Standardised inoculum across all agars consisted of diluted pure cultures of diverse *E. coli* strains. Key: MHA -Mueller-Hinton agar, MAC -MacConkey agar, Brill -Brilliance^™^ *E. coli* agar, Chrom -CHROMagar^™^ ECC agar.

All *E. coli* strains susceptible to ampicillin, tetracycline, ciprofloxacin and ceftiofur did not grow on agars with the corresponding incorporated antimicrobial (at any concentration). However, *E. coli* strains susceptible to gentamicin grew on MAC (2 and 4 μg/mL), Brilliance (2 μg/mL) and CHROMagar (2 μg/mL) agars incorporated with gentamicin. All *E. coli* strains resistant to ampicillin, tetracycline and ciprofloxacin grew on all agars with the corresponding incorporated antimicrobial (at any concentration). SA1001 was the only gentamicin-resistant *E. coli* strain that grew on all agars incorporated with gentamicin at all concentrations while growth of SA44 (also resistant to gentamicin) was not observed on MHA incorporated with 8 and 16 μg/mL of gentamicin. Ceftiofur-resistant *E. coli* strains grew on MAC agar when incorporated with ceftiofur (at any concentration). In contrast, growth was inconsistent on Brilliance and CHROMagar agars when incorporated ceftiofur concentrations climbed above 1 μg/mL (Fig. S2).

Separate linear models were constructed for each antimicrobial used. As with agar without antimicrobials, Brilliance and CHROMagar agars performed consistently better than MAC agar (Fig. 2). This includes Brilliance and CHROMagar agars incorporated with ceftiofur which was superior to MAC agar incorporated with ceftiofur (Fig. 2). Antimicrobial concentration was found to have a significant effect for all antimicrobials tested (P<0.001). Agar had a significant effect on all antimicrobials except tetracycline (P<0.05) and strain had significant effects on all except tetracycline and gentamicin (P<0.01). Significant interaction effects between strain and agar were found for tetracycline, gentamicin and ceftiofur (P<0.05), between agar and antimicrobial concentration for tetracycline (P<0.01) and ceftiofur (P<0.001), between strain and antimicrobial concentration for tetracycline and gentamicin (P<0.05) and between all three factors for gentamicin (P<0.01).

**FIG 2.**
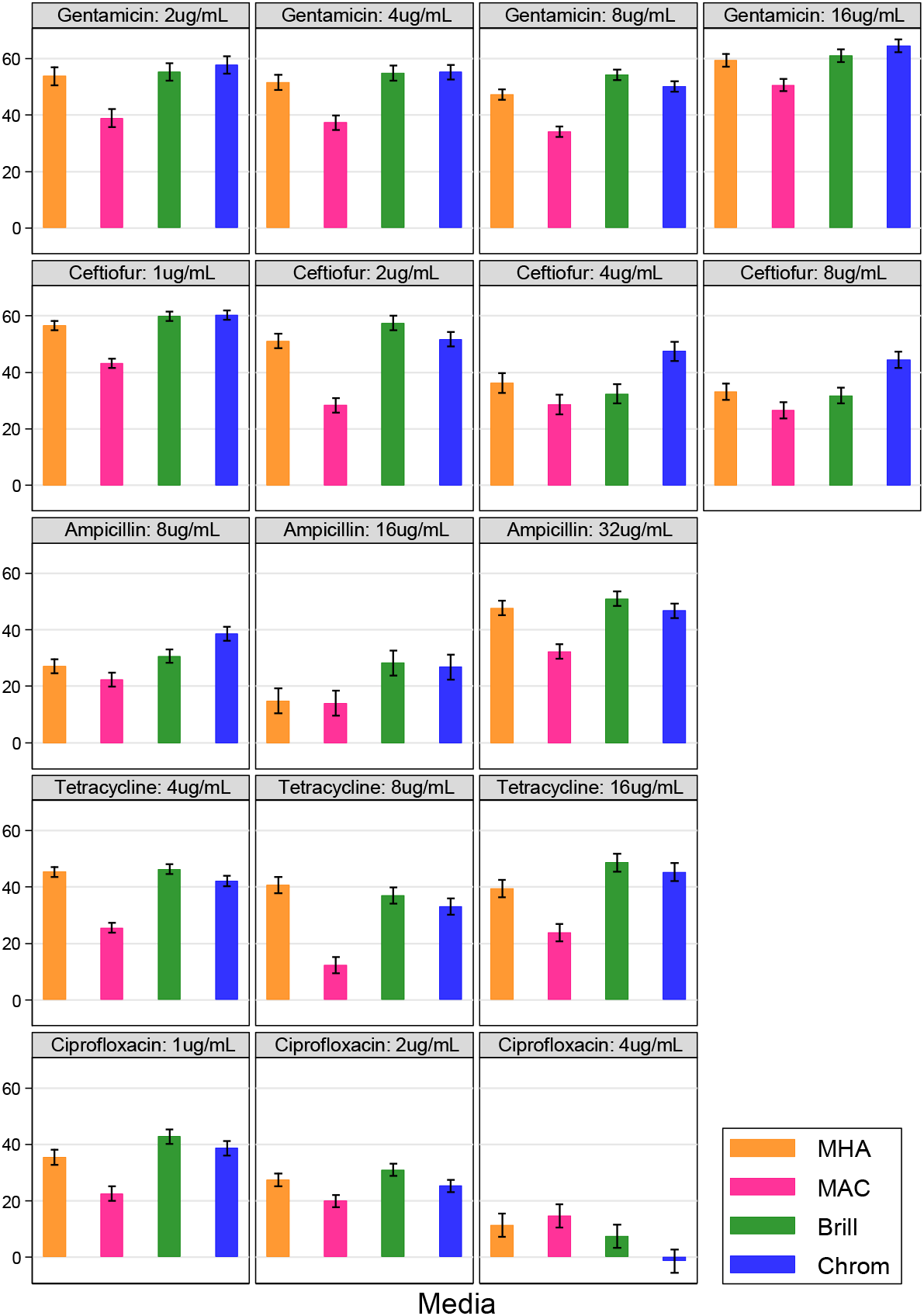
Comparisons of *E. coli* growth performance (mean colony counts per plate ± se) on three *E. coli* selective agars and Mueller-Hinton agar each incorporated with ampicillin, tetracycline, gentamicin, ciprofloxacin or ceftiofur at three or four concentrations (total number of plates = 424). Standardised inoculum across all agars consisted of diluted pure cultures of diverse *E. coli* strains resistant to each antimicrobial. Key: MHA -Mueller-Hinton agar, MAC -MacConkey agar, Brill -Brilliance^™^ *E. coli* agar, Chrom -CHROMagar^™^ ECC agar.

Finally, the three *E. coli* selective agars with and without incorporation of antimicrobials was further tested using homogenised bovine faecal samples (with and without spiking of two fluoroquinolone [FQ]-resistant *E. coli* strains: Table S2). For agars without antimicrobials, the order of superiority was CHROMagar (mean of 35.6 colonies per plate), Brilliance (mean of 34.2 colonies per plate) and MAC agars (mean of 29.1 colonies per plate) (Table 1). For agars incorporated with ciprofloxacin, growth of FQ-resistant *E. coli* strains was observed on all agars regardless of bacterial concentration and the order of superiority was Brilliance (mean of 32.8 colonies per plate), CHROMagar (mean of 28.3 colonies per plate) and MAC (mean of 22.8 colonies per plate) agars (Table 1). In this model, agar (P<0.001), strain (P<0.05), bacterial concentration (P<0.001) and interactions between agar and bacterial concentration had significant effects (P<0.001).

**Table 1.** Comparisons of *E. coli* and fluoroquinolone-resistant *E. coli* growth performance (mean colony counts per plate) on *E. coli* selective agar with and without incorporation of ciprofloxacin (total number of plates = 288). Standardised inoculum consisted of homogenised bovine faecal samples with and without spiking of fluoroquinolone-resistant *E. coli* strains. Key: MAC - MacConkey agar, Brilliance - Brilliance^™^ *E. coli* agar, CHROMagar - CHROMagar^™^ ECC agar.

| Homogenised bovine<br>faecal samples | MAC |  | Brilliance |  | CHROMagar |  |
| --- | --- | --- | --- | --- | --- | --- |
|  | Without<br>antimicrobials | With<br>ciprofloxacin | Without<br>antimicrobials | With<br>ciprofloxacin | Without<br>antimicrobials | With<br>ciprofloxacin |
| **Without**<br>**spiking** | 29.1 | - | 34.2 | - | 35.6 | - |
| **Spiked with**<br>**fluoroquinolone-**<br>**resistant *E. coli* strains** | - | 22.8 | - | 28.3 | - | 32.8 |

### Experiment B: Comparison of ESC-resistant *E. coli* growth on commercial ESC-resistant *E. coli s*elective agars

Two ESC-resistant *E. coli* selective agars (Brilliance ESBL and CHROMagar ESBL agars) were compared for the ability to support growth of diverse ESC-resistant *E. coli* strains (Table S2). The non-selective MHA without antimicrobials was used as a control agar. All ESC-resistant *E. coli* strains grew on all agars with the exception of SA27 which did not grow on Brilliance ESBL agar. CHROMagar ESBL agar (mean of 48.24 colonies per plate) best supported growth followed by MHA (mean of 42.72 colonies per plate) and Brilliance ESBL agar (mean of 28.74 colonies per plate) (Fig. 3) and this order of superiority was also observed on each ESC-resistant *E. coli* strain (Fig. S3). In this model, all factors and their associated interactions (P<0.001) had significant effects on colony counts.

**FIG 3.**
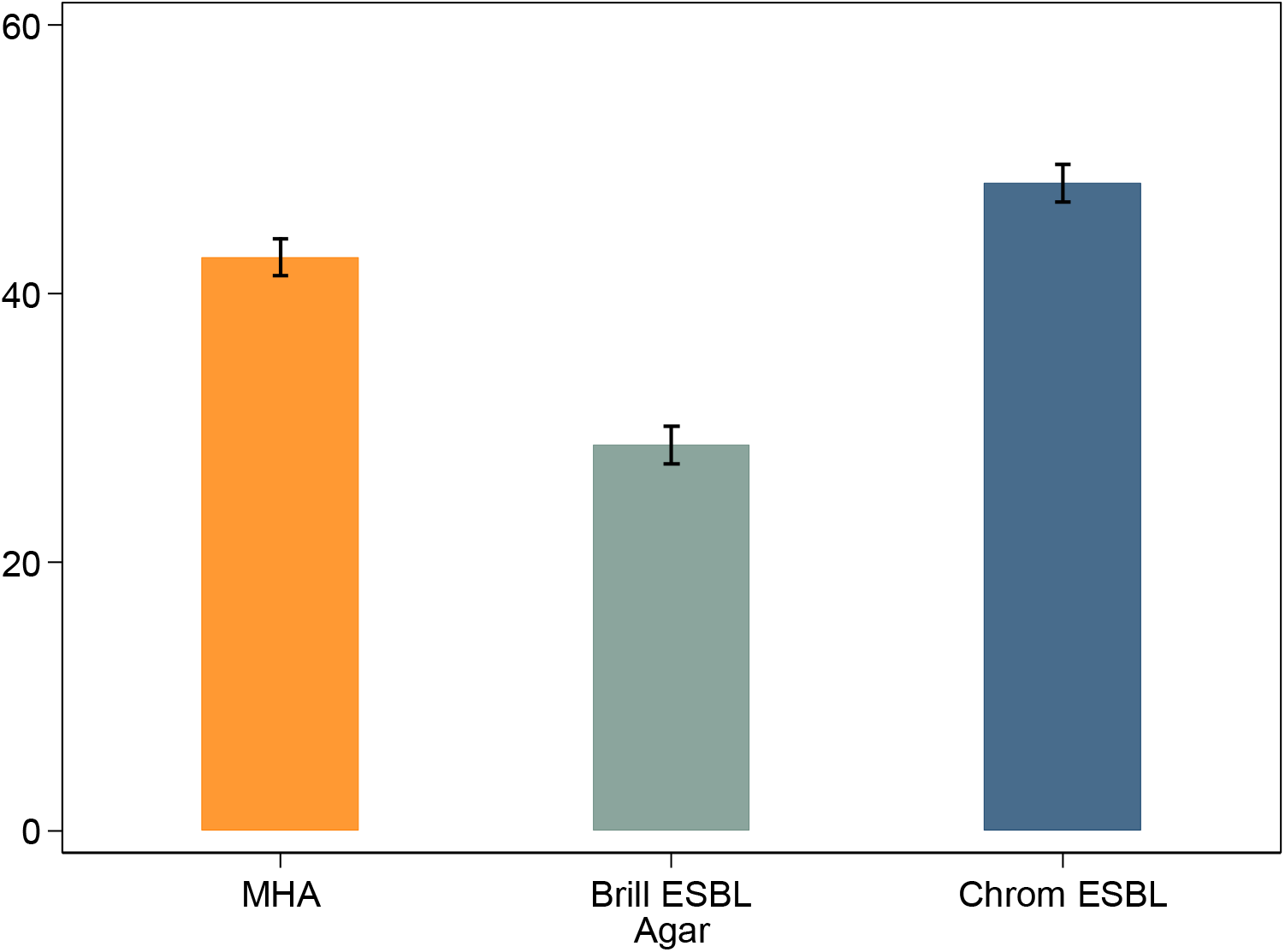
Comparisons of ESC-resistant *E. coli* growth performance (mean colony counts per plate ± se) on two ESC-resistant *E. coli* selective agars with Mueller-Hinton agar (without antimicrobials) present as a control agar (total number of plates = 150). Standardised inoculum across all agars consisted of diluted pure cultures of diverse ESC-resistant *E. coli* strains. Key: MHA -Mueller-Hinton agar, Brill ESBL -Brilliance™ ESBL agar, Chrom ESBL -CHROMagar^™^ ESBL agar.

Finally, Brilliance ESBL and CHROMagar ESBL agars were further tested using homogenised bovine faecal samples spiked with ten ESC-resistant *E. coli* strains (Table S2). Brilliance ESBL agar (mean of 24.3 colonies per plate) was found to be superior to CHROMagar ESBL agar (mean of 14.9 colonies per plate) (Fig. 4) with the same superiority order observed on each ESC-resistant *E. coli* strain (Fig. S4). The only exception was SA27 which did not grow on Brilliance ESBL agar regardless of bacterial concentration. All factors including associated interactions had significant effects (P<0.001) on colony counts.

**FIG 4.**
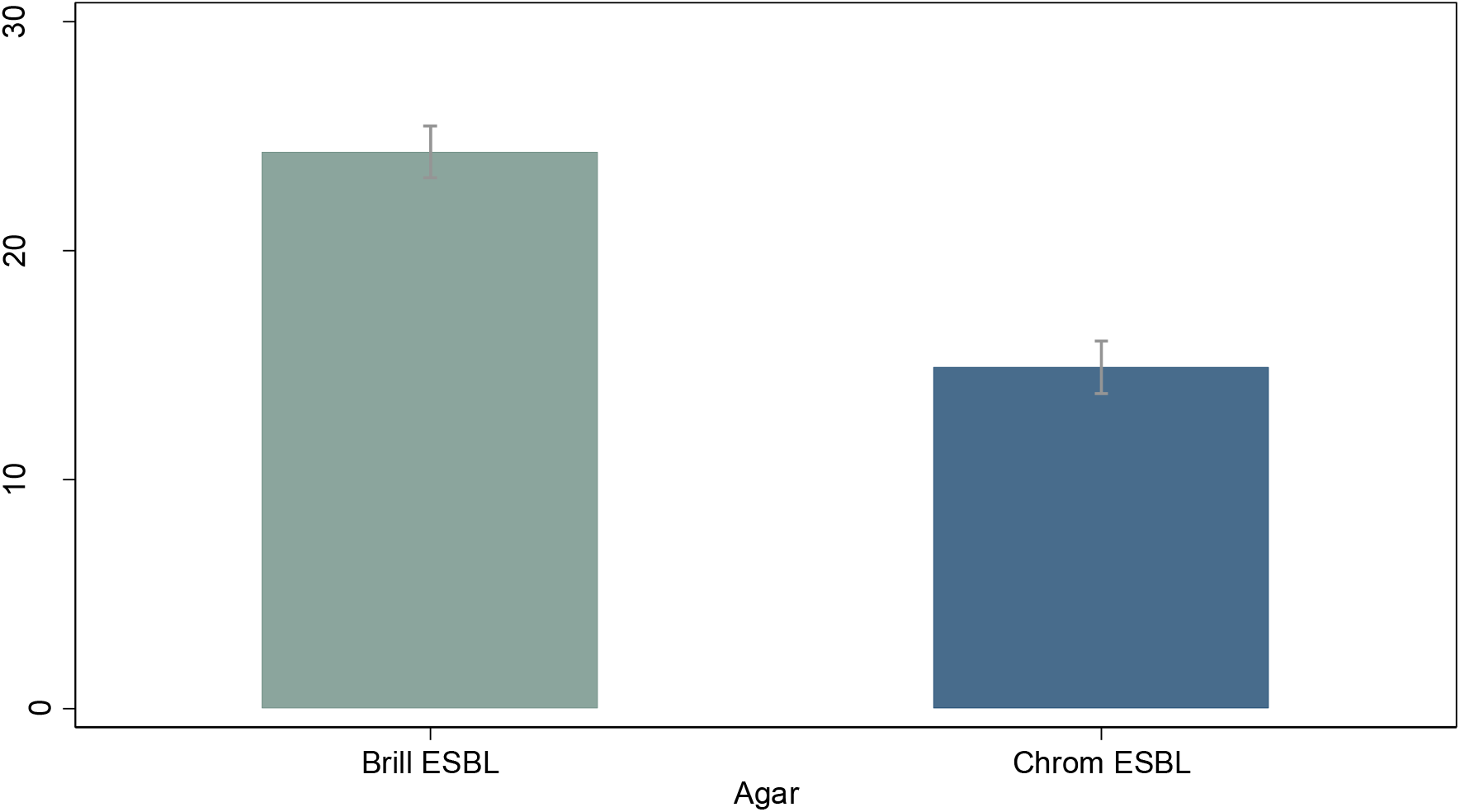
Comparisons of ESC-resistant *E. coli* growth performance (mean colony counts per plate ± se) on two ESC-resistant *E. coli* selective agars (total number of plates = 180). Standardised inoculum across all agars consisted of homogenised bovine faecal samples spiked with diverse ESC-resistant *E. coli* strains. Key: Brill ESBL -Brilliance^™^ ESBL agar, Chrom ESBL -CHROMagar^™^ ESBL agar.

## DISCUSSION

AMR surveillance in livestock and food products is a critical tool for progressive antimicrobial stewardship, prevention of AMR spread and the preservation of effective antimicrobials. Through the combination of high-throughput robotics with selective agar incorporated with the antimicrobial of interest, it is possible to quantify carriage levels and prevalence of resistance. With *E. coli* being used as a common indicator bacterium in AMR surveillance systems (6), this study aimed to identify the optimal selective agar and antimicrobial concentrations for quantifying populations of resistant *E. coli* for AMR surveillance in livestock.

In this study, three selective agars were tested (MAC, Brilliance and CHROMagar agars). Despite the presence of other experimental factors and interactions significantly affecting colony counts, both Brilliance and CHROMagar agars were comparable in performance and consistently superior to MAC agar in all situations (pure cultures, faecal samples and faecal samples spiked with FQ-resistant *E. coli* strains) as demonstrated through a higher number of *E. coli* colonies. The superior growth performance on Brilliance and CHROMagar agars can be attributed to the basic function and design of the agar. Both selective agars were specifically formulated for growing coliform bacteria and while the exact ingredients within the selective mix of both agars are undisclosed by the manufacturer, there may be components that provide specific growth support towards coliform bacteria including *E. coli*. In contrast, the consistently inferior performance by MAC agar could be attributed to its components that indiscriminately select for Gram-negative bacteria. Unlike Brilliance and CHROMagar agars, MAC agar possesses bile salts as its selective component to suppress Gram-positive bacteria growth by induction of DNA damage (20). However, it is also likely that this bile salt mechanism also indirectly exerts a suppressive effect on *E. coli* growth due to *E. coli* constantly having to express genes that reduces growth rate in order to repair any DNA damage (21). Therefore, the indiscriminate selection combined with the suppressive effect of bile salts in MAC agar presents a more stressful environment for *E. coli* resulting in an inferior performance which is confirmed in Fig 1. Additionally, this consistent performance of Brilliance and CHROMagar agars also demonstrated that the capability of both agars in supporting susceptible and FQ-resistant *E. coli* growth for detection and quantification was not impeded by the co-presence of faecal microflora.

Ampicillin, tetracycline, gentamicin, ciprofloxacin and ceftiofur were incorporated into each agar to identify the best concentration for growing the corresponding resistant *E. coli* for quantification. MAC agar consistently supported less growth regardless of antimicrobial and concentration and thus is not considered appropriate for quantitative AMR surveillance. Only ampicillin, tetracycline and ciprofloxacin were found to be suitable for incorporation into Brilliance and CHROMagar agars at all defined concentrations with growth of all resistant strains observed. In contrast, gentamicin was only suitable for incorporation into Brilliance and CHROMagar agars at 8 and 16 μg/mL as growth of susceptible strains were observed at lower concentrations. A higher number of susceptible strains grew on CHROMagar agar than Brilliance agar which suggests a higher level of suppression of gentamicin activity with the former. Currently, it is difficult to ascertain the mechanism by which this suppression occurs, although one possibility could be due to the significant three-way interaction between all factors.

A significant interaction between agars and ceftiofur was identified which, given the unexpected growth inhibition of some ceftiofur-resistant *E. coli* strains at higher ceftiofur concentrations, indicate a likely amplification of ceftiofur activity when incorporated into each agar. It is also possible that this amplification also extends to 1 μg/mL despite all ceftiofur-resistant *E. coli* strains growing at this concentration. With the lack of information in the current literature pertaining to interactions between agar and ceftiofur, further investigation is needed to explain this phenomenon. Nonetheless, this indicates that ceftiofur is not suitable for incorporation into Brilliance and CHROMagar agars and we suggest that either ESC-resistant *E. coli* selective agar such as CHROMagar ESBL agar be used for quantitative AMR surveillance of ESC-resistant *E. coli* or further investigation into the viability of using other third-generation cephalosporin antimicrobials such as cefotaxime or ceftriaxone for incorporation into selective agar.

Finally, both Brilliance ESBL and CHROMagar ESBL agars had unique advantages. While Brilliance ESBL agar was superior in supporting growth of ESC-resistant *E. coli* strains from spiked homogenised faecal samples, CHROMagar ESBL agar was able to support a wider diversity of ESC-resistant *E. coli* strain. This was evident from the absence of SA27 growth on Brilliance agar as opposed to its presence of growth on CHROMagar agar regardless if it was from a pure culture or spiked homogenised faecal sample which also serves to demonstrate that the interference of SA27 (and thus ESC-resistant *E. coli*) growth on both agars may likely be due to interactions between strain and agar rather than the co-presence of faecal microflora. Nevertheless, the capability of CHROMagar ESBL agar to capture a wider diversity of ESC-resistant *E. coli* makes it better suited for AMR surveillance than Brilliance ESBL agar as it would increase the probability of detecting ESC-resistant *E. coli*.

The reason for growth variation between strains was not clear as it was not the principal feature being evaluated. Most data in the current literature focuses on growth rate of *E. coli* strains under specific environmental conditions but none have evaluated possible factors influencing growth rates between *E. coli* strains (22-24). Significant interactions between strain with agar or antimicrobial are one such factor affecting growth rate but given the uniformity in performance across all agars with and without incorporation of antimicrobials, it suggests that this influence towards growth was minimal and not enough to affect the performance outcome of each agar.

This study represents the first step towards establishing an enhanced AMR surveillance approach for assessing AMR in livestock and food products. As opposed to the established approach of AMR surveillance, this enhanced approach is both qualitative and quantitative in nature and is built on the capacity to rapidly identify *E. coli* colonies on agars for colony enumeration. When combined with robotics, it provides exciting opportunities for up-scaling based on programming and machine learning pathways to allow the identification of *E. coli* colonies based on colony colour for enumeration with reduced human input and potentially greater accuracy. The practical ramifications for this are that more accurate information can be obtained from a greater number of samples that increases the sensitivity of detecting a given phenotype across a population of animals and herds. It is an especially relevant technique for early detection of resistance to critically important antimicrobials (CIAs) since it cannot be assumed that either the level of colonisation is uniform across animals or herds (25), or that the phenotypes of interest are present at a high enough concentration to be found by traditional AST means. Moreover, any positive colonies detected can be preserved for genomic interrogation to understand their ecological origins as demonstrated in studies of human-wildlife-livestock transmission (26).

Based on this study, we recommend the use of Brilliance and CHROMagar agars with and without incorporation of antimicrobials as well as CHROMagar ESBL agar in combination with robotics to evaluate the feasibility of this enhanced approach. Additionally, this enhanced approach also has promising applications within food, clinical and public health settings through large-scale qualitative and quantitative AMR surveillance of critically important antimicrobial-resistant bacteria to support infection control and evaluation of the effectiveness of antimicrobial stewardship (27).

## MATERIALS AND METHODS

All agars used in this study were commercially available and were used as directed by the manufacturer with the exception of the incorporation of additional antimicrobials as demanded by study design.

### Experiment A: Comparison of *E. coli* selective agars with and without incorporation of antimicrobials

The performance of growing *E. coli* on three *E. coli* selective agars and a fourth non-selective control agar with and without incorporation of antimicrobials were compared using pure cultures of diverse *E. coli* strains. All agars without antimicrobials were purchased directly from suppliers. The three selective agars used were MAC (Edwards Group), Brilliance (Thermo Fisher Scientific) and CHROMagar (MicroMedia, Edwards Group) agars. MHA (Edwards Group) was used as the fourth agar and was chosen for comparison due to its status as the gold standard non-selective agar for routine AST (28). The same four agars incorporated with antimicrobials were prepared in-house using the agar dilution technique as per manufacturer instructions. Both Mueller-Hinton broth powder (Oxoid, Thermo Fisher Scientific) and agar No. 1 powder (Oxoid, Thermo Fisher Scientific) were used to prepare MHA. MacConkey No. 3 powder (Oxoid, Thermo Fisher Scientific) was used to prepare MAC agar. Brilliance agar was prepared using Brilliance™ *E. coli*/coliform selective medium powder (Oxoid, Thermo Fisher Scientific) while *E. coli*-coliforms chromogenic medium (Conda, Edwards Group) was used to prepare CHROMagar agar. The antimicrobials selected for incorporation into agars were ampicillin, tetracycline, gentamicin, ciprofloxacin and ceftiofur (from the penicillin, tetracycline, aminoglycoside, FQ and third-generation cephalosporin families respectively). These were included due to their importance in the livestock and public health sectors (particularly ciprofloxacin and ceftiofur which are CIAs for human medicine) and thus often included in AMR surveillance programs involving livestock and food products (3, 5, 29-32). All antimicrobial stocks were prepared using antimicrobial powders (Sigma-Aldrich) and stored following Clinical and Laboratory Standards Institute (CLSI) guidelines (28). All stocks were used within the shelf life detailed by the manufacturer. Prior to pouring into petri dishes, antimicrobials were added to sterilised agars after being cooled in a 60 °C water bath, to obtain specific concentrations for each respective antimicrobial. Three concentrations were chosen for ampicillin (8, 16 and 32 μg/mL), tetracycline (4, 8 and 16 μg/mL) and ciprofloxacin (1, 2 and 4 μg/mL) while four were chosen for gentamicin (2, 4, 8 and 16 μg/mL) and ceftiofur (1, 2, 4 and 8 μg/mL). All concentrations were chosen to cover the clinical breakpoints for *E. coli* as listed by CLSI with the epidemiological cut-off points (ECOFF) listed by the European Committee of Antimicrobial Susceptibility Testing (EUCAST) covered for ampicillin, tetracycline, gentamicin and ceftiofur (33). After the addition of antimicrobials, 20 mL of the agar mixture was poured into 90 mm diameter circular petri dishes and left to solidify under a laminar flow hood. All agars incorporated with antimicrobials were stored in the dark at 4 °C and used within two weeks of preparation.

Eight *E. coli* strains were chosen with ATCC 25922 included as the quality control strain while the remaining seven strains were *E. coli* isolated from different animal species. SA44 was isolated from pigs (34), SA1000, SA1001 and SA1002 were isolated from Australian Silver Gulls (26), and SA1003, SA1004 and SA1005 were archival in-house strains (Table S1). The rationale for selection of these strains was to achieve diversity in origin to capture variations potentially present in wild type populations of *E. coli*. Prior to the commencement of the experiment, minimal inhibitory concentration (MIC) testing using the microbroth dilution method was performed on all *E. coli* strains according to CLSI guidelines to confirm the resistance profile of each strain (28) (Table S1). All growth observations on agars incorporated with antimicrobials were compared to the resistance profile shown for each strain. After overnight growth on Columbia sheep blood agar (Edwards Group), a suspension of each *E. coli* strain meeting the 0.5 McFarland standard was prepared using a nephelometer (Sensititre). Each standardised inoculum underwent 10-fold serial dilution to 10^™5^ in sterile 1 x phosphate buffered saline. Inoculation was performed by dispensing 80 μL of the 10^™5^ inoculum onto agar without antimicrobials and spread evenly across the agar surface using a sterile loop. Inoculation on agars without antimicrobials was repeated for a total of five replicates per combination of agar and strain while inoculation on agars incorporated with antimicrobials was repeated for a total of two replicates per combination of agar, strain and antimicrobial concentration. All agars were incubated between 16 to 20 hours at 37 °C.

Presumptive identification of *E. coli* on Brilliance and CHROMagar agars was performed based on colony colour as detailed by the manufacturer. For MAC agar, pink colonies were presumed to be *E. coli* due to most *E. coli* strains being known to be lactose fermenters. Being a non-selective agar, *E. coli* colonies on MHA appear colourless.

Homogenised bovine faecal samples were used as field samples to verify the performance for growing *E. coli* on the same three *E. coli* selective agars without antimicrobials. All agars without antimicrobials were purchased from the same suppliers described above. Twenty bovine faecal samples from the Murdoch University farm were sampled. All faecal samples were collected from fresh faecal piles and processed on the same day of collection. Approximately 2 g of each faecal sample was homogenised for 30 seconds in 18 mL of sterile 1x phosphate buffered saline (PBS) using a BagMixer^®^ 400 P laboratory blender (Interscience, Edwards Group). This was repeated two more times to obtain a total of three replicates per sample. The homogenised mixture of each replicate underwent a 10-fold serial dilution and 80 μL of each dilution factor at 10^™1^ was inoculated onto each agar and spread evenly across the agar surface using a sterile loop. The procedure for agar incubation and presumptive identification of *E. coli* on agar were the same described above for agars without antimicrobials.

Homogenised bovine faecal samples spiked with FQ-resistant *E. coli* strains was used as field samples to further evaluate the performance for growing FQ-resistant *E. coli* on the same three *E. coli* selective agars incorporated with 4 μg/mL of ciprofloxacin (Table S2). All agars were prepared in the same manner described above for agars incorporated with antimicrobials.

A ST131 and ST744 *E. coli* strain isolated from Australian Silver Gulls was chosen for inoculation into faecal samples due to their ubiquity as FQ-resistant *E. coli* strains internationally in both humans and animals (26, 35-38). The first ten bovine faecal samples used previously were chosen for pooling. Each pooled sample consists of five individual samples to form a total of two pooled samples. For each pooled sample, approximately 2 g of each individual faecal sample (total of approximately 10 g) was homogenised for 30 seconds in 90 mL of sterile 1× phosphate buffered saline (PBS) using a BagMixer^®^ 400 P laboratory blender (Interscience, Edwards Group). This was repeated two more times to obtain a total of three replicates per pool sample. After overnight growth on Columbia sheep blood agar (Edwards Group), a suspension of each *E. coli* strain meeting the 0.5 McFarland standard was prepared using a nephelometer (Sensititre) and inoculated into the homogenised mixture of each replicate to obtain bacterial concentrations of 10^3^, 10^5^ and 10^7^ colony forming units per gram (CFU/g). Mixtures containing 10^5^ and 10^7^ CFU/g were serially diluted to 10^™1^ and 10^™3^ dilution factor respectively. 80 μL of 10^3^, 10^5^ and 10^7^ at neat, 10^™1^ and 10^™3^ dilution factor respectively were inoculated onto each agar and spread evenly across the agar surface using a sterile loop. The procedure for agar incubation and presumptive identification of *E. coli* on agar were the same as described above for agars incorporated with antimicrobials. ATCC 25922 was also inoculated onto each agar as quality control.

### Experiment B: Comparison of ESC-resistant *E. coli* selective agars

The performance of growing ESC-resistant *E. coli* on two ESC-resistant *E. coli* selective agars were compared using pure cultures of diverse ESC-resistant *E. coli* strains (Table S2). All agars were purchased directly from suppliers. Brilliance ESBL (Thermo Fisher Scientific) and CHROMagar ESBL (MicroMedia, Edwards Group) agars were the two ESC-resistant *E. coli* selective agars while MHA (Thermo Fisher Scientific) was selected as the non-selective agar. The supplier for MHA in Experiment B differed from Experiment A, however the formulation of the agar was the same. Ten ESC-resistant *E. coli* strains were chosen with each strain harbouring a different gene conferring resistance to ESCs in order to encompass the wide genotypic variations present in ESC-resistant *E. coli* strains (Table S2). SA44 and SA1001 were the only two strains from Experiment A included in Experiment B (Table S2). Of the remaining eight strains, SA27 was isolated from pigs (35) while SA1074, SA1075, SA1076, SA1077, SA1078, SA1079 and SA1080 were isolated from Australian Silver Gulls (26) (Table S2). The procedure for culturing ESC-resistant *E. coli* strains, McFarland standard preparation, agar inoculation (including replicate numbers) and incubation, and presumptive identification of *E. coli* on MHA were the same as Experiment A using pure cultures of *E. coli* strains. Presumptive identification of ESC-resistant *E. coli* on Brilliance ESBL and CHROMagar ESBL agars were performed based on colony colour detailed by the manufacturer. ATCC 25922 was also inoculated onto each agar as quality control.

Homogenised bovine faecal samples spiked with ESC-resistant *E. coli* strains was used as field samples to verify the performance for growing ESC-resistant *E. coli* on the same two ESC-resistant *E. coli* selective agars (Table S2). Ten ESC-resistant *E. coli* strains were also chosen with nine strains, SA27, SA44, SA1001, SA1074, SA1075, SA1076, SA1077, SA1079 and SA1080, being the same strains described above while the last strain, SA1083, was another strain previously isolated from Australian Silver Gulls (26) (Table S2). The first five bovine faecal samples used previously in Experiment A were chosen for pooling. The procedure for pooling, strain inoculation into homogenised faecal mixture, agar inoculation (including replicate numbers) and incubation were the same as Experiment A when using homogenised bovine faecal samples spiked with FQ-resistant *E. coli* strains on agars incorporated with ciprofloxacin with the exception that only Brilliance ESBL and CHROMagar ESBL agars were used. Presumptive identification of ESC-resistant *E. coli* on agar was the same as described above. ATCC 25922 was also inoculated onto each agar as quality control.

### Statistical analysis

Statistical analysis used the linear model framework in Stata version 16.0 (Stata Corporation, TX, USA). All analyses were fixed effect models with the count of *E. coli* colonies on each plate as the outcome with results expressed (in text and figures) as the mean effect of each level of the factor of interest (eg. agar type or bacterial strain) and adjusted for other terms in the model. For experiments based on pure cultures of *E. coli* strains, a model was constructed for agars without antimicrobials, and one model for each antimicrobial when incorporated into agars. In the latter case, only *E. coli* strains resistant to the antimicrobial being evaluated was included in the linear model. For experiments based on faecal samples spiked with a mixture of *E. coli* strains, the analysis was similar although the factor representing bacterial strain was not.

## Acknowledgements

This project was supported by the Australian Department of Agriculture, Water and the Environment. We thank Professor John Turnidge for critical review of the manuscript.

## Ethical approval

Not required

## REFERENCES

1. World Health Organization. 2014. Antimicrobial resistance: Global report on surveillance. World Health Organization, Switzerland.

2. Aarestrup FM, Wegener HC, Collignon P. 2008. Resistance in bacteria of the food chain: epidemiology and control strategies. Expert Rev Anti Infect Ther 6:733–750.

3. Abraham S, O’Dea M, Sahibzada S, Hewson K, Pavic A, Veltman T, Abraham R, Harris T, Trott DJ, Jordan D. 2019. Escherichia coli and Salmonella spp. isolated from Australian meat chickens remain susceptible to critically important antimicrobial agents. PLoS One 14.

4. Mukerji S, O’Dea M, Barton M, Kirkwood R, Lee T, Abraham S. 2017. Development and transmission of antimicrobial resistance among Gram-negative bacteria in animals and their public health impact. Essays Biochem 61:23–35.

5. Nekouei O, Checkley S, Waldner C, Smith BA, Invik J, Carson C, Avery B, Sanchez J, Gow S. 2018. Exposure to antimicrobial-resistant *Escherichia coli* through the consumption of ground beef in Western Canada. Int J Food Microbiol 272:41–48.

6. World Health Organization. 2021. Global tricycle surveillance ESBL *E. coli*. World Health Organization, Switzerland.

7. European Food Safety Authority, Aerts M, Battisti A, Hendriksen R, Kempf I, Teale C, Tenhagen BA, Veldman K, Wasyl D, Guerra B, Liebana E, Thomas-Lopez D, Beloeil PA. 2019. Technical specifications on harmonised monitoring of antimicrobial resistance in zoonotic and indicator bacteria from food-producing animals and food. EFSA J 17.

8. Humphries RM, Ambler J, Mitchell SL, Castanheira M, Dingle T, Hindler JA, Koeth L, Sei K, CLSI Methods Development and Standardization Working Group of the Subcommittee on Antimicrobial Susceptibility Testing. 2018. CLSI methods development and standardization working group best practices for evaluation of antimicrobial susceptibility tests. J Clin Microbiol 56.

9. DANMAP. 2017. DANMAP 2016 - Use of antimicrobial agents and occurrence of antimicrobial resistance in bacteria from food animals, food and humans in Denmark. DANMAP, Denmark.

10. Truswell A, Abraham R, O’Dea M, Lee ZZ, Lee T, Laird T, Blinco J, Kaplan S, Turnidge J, Trott DJ, Jordan D, Abraham S. 2021. Robotic Antimicrobial Susceptibility Platform (RASP): a next-generation approach to One Health surveillance of antimicrobial resistance. J Antimicrob Chemother doi:10.1093/jac/dkab107.

11. Perry JD, Freydiere AM. 2007. The application of chromogenic media in clinical microbiology. J Appl Microbiol 103:2046–2055.

12. Grohs P, Tillecovidin B, Caumont-Prim A, Carbonnelle E, Day N, Podglajen I, Gutmann L. 2013. Comparison of five media for detection of extended-spectrum bea-lactamase by use of the Wasp instrument for automated specimen processing. J Clin Microbiol 51:2713–2716.

13. Huang TD, Bogaerts P, Berhin C, Guisset A, Glupczynski Y. 2010. Evaluation of Brilliance ESBL agar, a novel chromogenic medium for detection of extended-spectrum-beta-lactamase-producing *Enterobacteriaceae*. J Clin Microbiol 48:2091–2096.

14. Pultz MJ, Nerandzic MM, Stiefel U, Donskey CJ. 2008. Emergence and acquisition of fluoroquinolone-resistant Gram-negative bacilli in the intestinal tracts of mice treated with fluoroquinolone antimicrobial agents. Antimicrob Agents Chemother 52:3457–3460.

15. Blaser J, Luthy R. 1988. Comparative study on antagonistic effects of low pH and cation supplementation on in-vitro activity of quinolones and aminoglycosides against *Pseudomonas aeruginosa*. J Antimicrob Chemother 22:15–22.

16. D’Amato RF, Thornsberry C, Baker CN, Kirven LA. 1975. Effect of calcium and magnesium ions on the susceptibility of *Pseudomonas* species to tetracycline, gentamicin, polymyxin B, and carbenicillin. Antimicrob Agents Chemother 7:596–600.

17. Fass RJ, Barnishan J. 1979. Effect of divalent cation concentrations on the antibiotic susceptibilities of nonfermenters other than *Pseudomonas aeruginosa*. Antimicrob Agents Chemother 16:434–438.

18. Girardello R, Bispo PJM, Yamanaka TM, Gales AC. 2012. Cation concentration variability of four distinct Mueller-Hinton agar brands influences polymyxin B susceptibility results. J Clin Microbiol 50:2414–2418.

19. Marshall AJ, Piddock LJ. 1994. Interaction of divalent cations, quinolones and bacteria. J Antimicrob Chemother 34:465–483.

20. Begley M, Gahan CG, Hill C. 2005. The interaction between bacteria and bile. FEMS Microbiol Rev 29:625–651.

21. Merritt ME, Donaldson JR. 2009. Effect of bile salts on the DNA and membrane integrity of enteric bacteria. J Med Microbiol 58:1533–1541.

22. Haberbeck LU, Oliveira RC, Vivijs B, Wenseleers T, Aertsen A, Michiels C, Geeraerd AH. 2015. Variability in growth/no growth boundaries of 188 different Escherichia coli strains reveals that approximately 75% have a higher growth probability under low pH conditions than E. coli O157:H7 strain ATCC 43888. Food Microbiol 45:222–30.

23. Smirnova GV, Oktyabrsky ON. 2018. Relationship between Escherichia coli growth rate and bacterial susceptibility to ciprofloxacin. FEMS Microbiol Lett 365.

24. Thakur CS, Brown ME, Sama JN, Jackson ME, Dayie TK. 2010. Growth of wildtype and mutant E. coli strains in minimal media for optimal production of nucleic acids for preparing labeled nucleotides. Appl Microbiol Biotechnol 88:771–9.

25. Persoons D, Bollaerts K, Smet A, Herman L, Heyndrickx M, Martel A, Butaye P, Catry B, Haesebrouck F, Dewulf J. 2011. The importance of sample size in the determination of a flock-level antimicrobial resistance profile for Escherichia coli in broilers. Microb Drug Resist 17:513–9.

26. Mukerji S, Stegger M, Truswell AV, Laird T, Jordan D, Abraham RJ, Harb A, Barton M, O’Dea M, Abraham S. 2019. Resistance to critically important antimicrobials in Australian silver gulls (*Chroicocephalus novaehollandiae*) and evidence of anthropogenic origins. J Antimicrob Chemother 74:2566–2574.

27. Perry JD. 2017. A decade of development of chromogenic culture media for clinical microbiology in an era of molecular diagnostics. Clin Microbiol Rev 30:449–479.

28. Clinical and Laboratory Standards Institute. 2018. M07-A11: Methods for dilution antimicrobial susceptibility tests for bacteria that grow aerobically; Approved Standard -Eleventh Edition. Clinical and Laboratory Standards Institute, USA.

29. Abraham S, O’Dea M, Page SW, Trott DJ. 2017. Current and future antimicrobial resistance issues for the Australian pig industry. Anim Prod Sci 57:2398–2407.

30. Kidsley AK, Abraham S, Bell JM, O’Dea M, Laird TJ, Jordan D, Mitchell P, McDevitt CA, Trott DJ. 2018. Antimicrobial susceptibility of *Escherichia coli* and *Salmonella* spp. isolates from healthy pigs in Australia: Results of a pilot national survey. Front Microbiol 9.

31. Moodley A, Guardabassi L. 2009. Transmission of IncN plasmids carrying *bla*_CTX-M-1_ between commensal *Escherichia coli* in pigs and farm workers. Antimicrob Agents Chemother 53:1709–1711.

32. van Breda LK, Dhungyel OP, Ward MP. 2018. Antibiotic resistant *Escherichia coli* in southeastern Australian pig herds and implications for surveillance. Zoonoses Public Health 65:e1–e7.

33. Clinical and Laboratory Standards Institute. 2018. M100: Performance Standards for Antimicrobial Susceptibility Testing - Twenty Eighth Edition. Clinical and Laboratory Standards Institute, USA.

34. Abraham S, Kirkwood RN, Laird T, Saputra S, Mitchell T, Singh M, Linn B, Abraham RJ, Pang S, Gordon DM, Trott DJ, O’Dea M. 2018. Dissemination and persistence of extended-spectrum cephalosporin-resistance encoding IncI1-blaCTXM-1 plasmid among *Escherichia coli* in pigs. ISME J 12:2352–2362.

35. Abraham S, Jordan D, Wong HS, Johnson JR, Toleman MA, Wakeham DL, Gordon DM, Turnidge JD, Mollinger JL, Gibson JS, Trott DJ. 2015. First detection of extended-spectrum cephalosporin- and fluoroquinolone-resistant *Escherichia coli* in Australian food-producing animals. J Glob Antimicrob Resist 3:273–277.

36. Garcia-Fernandez A, Villa L, Bibbolino G, Bressan A, Trancassini M, Pietropaolo V, Venditti M, Antonelli G, Carattoli A. 2020. Novel Insights and Features of the NDM-5-Producing Escherichia coli Sequence Type 167 High-Risk Clone. mSphere 5.

37. Gronthal T, Osterblad M, Eklund M, Jalava J, Nykasenoja S, Pekkanen K, Rantala M. 2018. Sharing more than friendship - transmission of NDM-5 ST167 and CTX-M-9 ST69 Escherichia coli between dogs and humans in a family, Finland, 2015. Euro Surveill 23.

38. Guenther S, Aschenbrenner K, Stamm I, Bethe A, Semmler T, Stubbe A, Stubbe M, Batsajkhan N, Glupczynski Y, Wieler LH, Ewers C. 2012. Comparable high rates of extended-spectrum-beta-lactamase-producing Escherichia coli in birds of prey from Germany and Mongolia. PLoS One 7:e53039.

